# CellView: Interactive exploration of high dimensional single cell RNA-seq data

**DOI:** 10.1101/123810

**Authors:** Mohan T. Bolisetty, Michael L. Stitzel, Paul Robson

## Abstract

Advances in high-throughput single cell transcriptomics technologies have revolutionized the study of complex tissues. It is now possible to measure gene expression across thousands of individual cells to define cell types and states. While powerful computational and statistical frameworks are emerging to analyze these complex datasets, a gap exists between this data and a biologist’s insight. The CellView web application fills this gap by providing easy and intuitive exploration of single cell transcriptome data.

---

Recent technological advances in single cell capture and nano-scale reactions have led to a major revolution in single cell transcriptomics^1,2,3^. Single cell datasets are analyzed using computational and statistical frameworks that enable feature (gene) selection, dimensionality reduction, clustering and differential gene expression. Multiple software packages exist that allow researchers well versed in computational analysis to perform this analysis^4–6^. However, identifying the exact parameters required for cell type identification is an iterative process greatly improved when informed by biology. In addition, interactive exploration of single cell datasets incorporating a biologist’s knowledge greatly improves data interpretation, yet often such experts do not have big data handling skills.

Advances in web application frameworks and visualization methods for dense datasets facilitate the development of interactive applications to allow easy and intuitive exploration of single cell data. Here, we introduce an R Shiny^7^ web application, CellView, that allows knowledge-based and hypothesis-driven exploration of processed single cell transcriptomic data. The input into CellView is an R dataset (.Rds) file with three pre-computed data frames containing expression, clustering, and gene symbol information. This file is agnostic of upstream computational approaches providing flexibility in algorithms used to calculate these data frames. This .Rds file can be shared with the end user, eliminating the need for hosting datasets, thereby decreasing the size of a virtual machine or cloud instance required to host and use CellView. Multiple tabs allow for easy access to the data and visualization of gene expression across and within clusters, aiding cell type identification.

To illustrate the utility and power of CellView, we generated and analyzed single cell transcriptome data from peripheral blood mononuclear cells (PBMCs) using the 10X Genomics Chromium^8^. As defined by the CellRanger^8^ pipeline, this data consisted of 6,554 single cells sequenced to 90.1% saturation with, on average, 824 genes and 2,077 molecules detected per cell. Dimensionality reduction using tSNE^9^ was applied to genes selected by normalized dispersion, and with clustering by DBSCAN^10^. CellView automatically determines cluster numbers, updates the user interface, and renders a 3D scatter plot displaying cells clustered in tSNE space (Fig 1b) from the uploaded .rds file.

**Figure 1:**
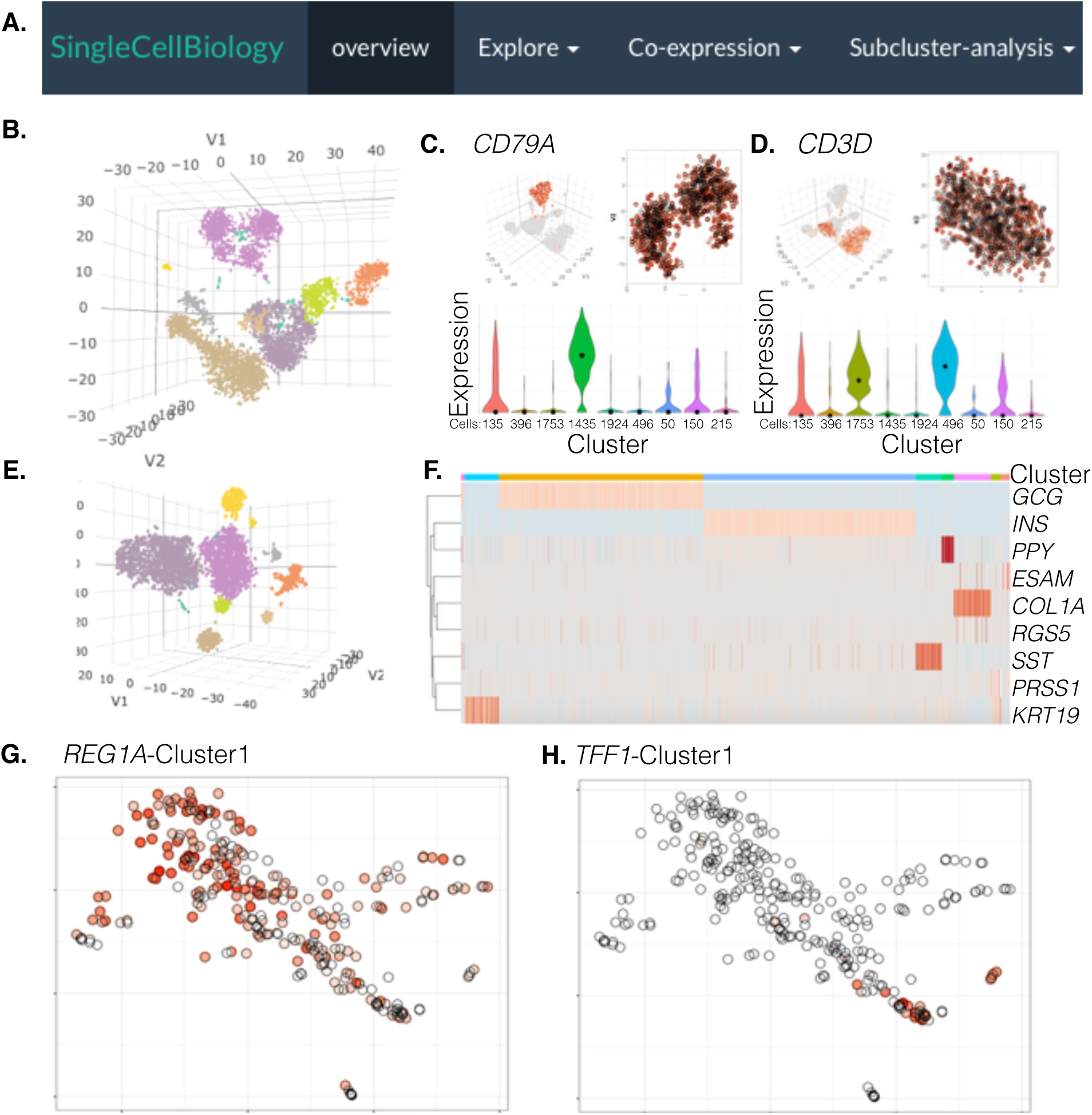
CellView enables cell type identification of clusters and discovery of novel cell states in PBMC and pancreatic islet datasets. **A.** CellView’s graphical user interface has 3 different features that enables exploration of single cell RNA-seq datasets. **B.** Upon PBMC data upload, a 3D plot of cells clustered in t-SNE space is displayed in ‘overview’. Expression patterns of marker genes such as **C.** *CD79A* and **D.** *CD3D* can be visualized in multiple panels under the ‘Explore’ module assisting in cell type identification and to discover further heterogeneity. **E.** 3D display of cell type clusters identified in human pancreatic islets. **F**. Analysis using the ‘Co-expression’ module of CellView with marker genes aids in the identification the major endocrine cell populations, alpha (cluster 2), beta (cluster 3), gamma (cluster 5), delta (cluster 4) along with exocrine cell types like ductal (cluster 1), stellate (cluster 6), acinar (cluster 7) and endothelial (cluster 8) cells. Cluster and gene specific views, **G.** *REG1A* and H. *TPP1* expression in the ductal cell cluster identifies cells in multiple states.

The ‘Explore’ tab provides cluster-centric exploration through three panel views. Panel 1 displays a 3D plot of a chosen gene’s expression across all cells. Panel 2 displays a 2D plot of the same gene’s expression across all cells in a single cluster, which users can select via drop-down list. Within Panel 2 users can download a .csv file of a gene-cell expression matrix by selecting cells with a square brush stroke. This provides convenient access to all genes expressed in a subset of cells. Panel 3 displays violin plots of the chosen gene’s cluster-specific expression and includes a total cell count for each cluster. *CD79A* (Fig 1c), a marker of B-cells, and *CD3D*, a marker of T-cells (Fig 1d), provide representative views of the ‘Explore’ tab.

The ‘Co-expression’ tab enables the generation of heatmaps to visualize expression of multiple genes either across all clusters, in the ‘AllClusters’ sub-menu, or on selected cells within a cluster, in the ‘Selected cells’ sub-menu (Fig 2a). The number of genes analyzed is only limited by legibility of the gene symbols in the resulting heatmap. This feature facilitates the use of known markers to empirically determine cell (sub)cluster identity.

**Figure 2:**
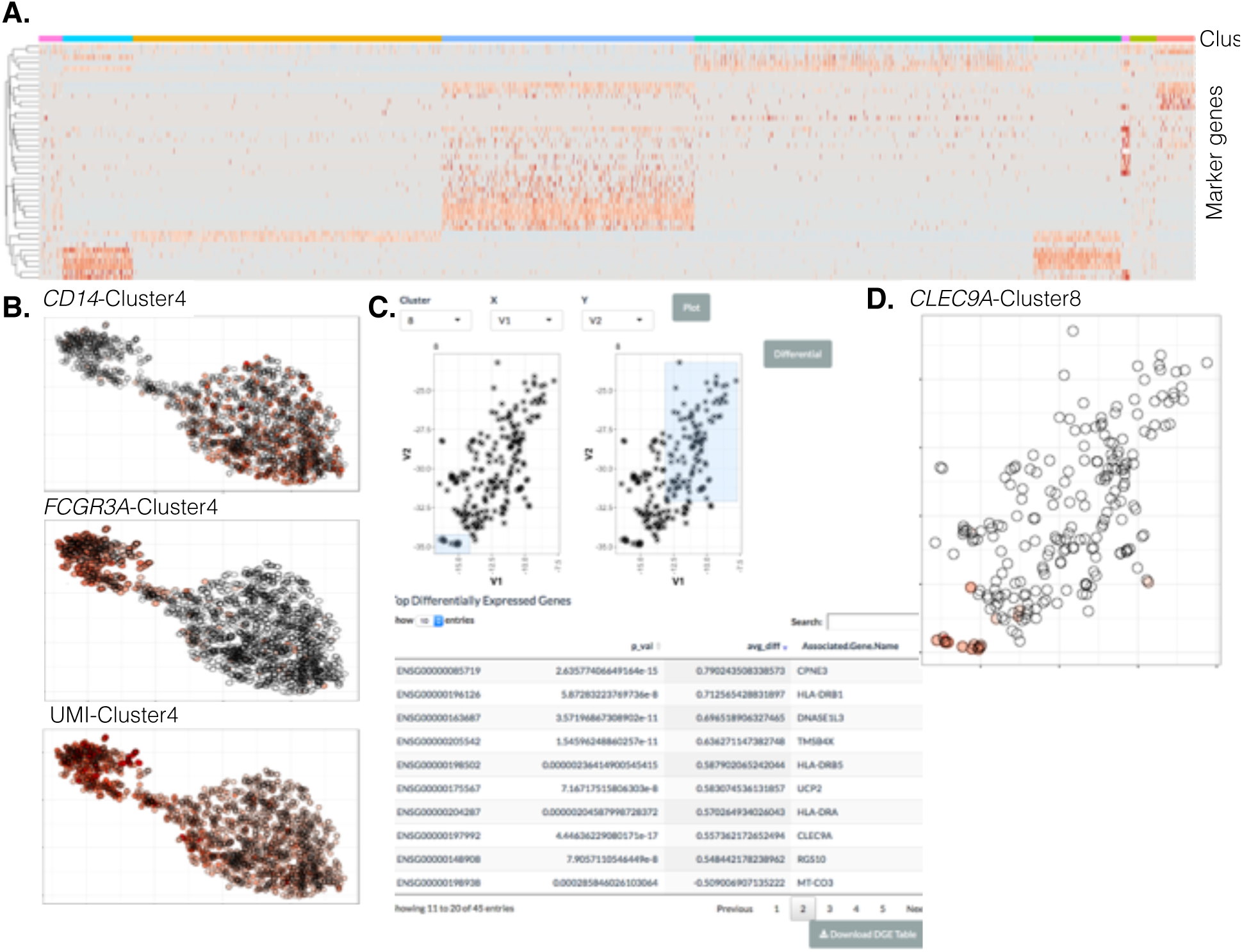
Investigating gene expression patterns across and within clusters with CellView identifies different cell type populations. **A.** Analysis using the co-expression module of CellView with various immune cell markers identifies the major subpopulations present in PBMCs. **B.** Exploring gene expression of *CD14* and *CD68* expression identifies a continuum from classical to non-classical monocytes that are also characterized by difference in absolute transcript counts (UMI). **C.** CellView allows for identification of sub-clusters by simple differential gene expression between groups of selected cells using a square brush stroke and displays a sortable and searchable table. **D.** Differential expression of two visually resolved populations in the dendritic cell cluster identifies less abundant CD141+ dendritic cells expressing the *CLEC9A* and *DNASE1L3* marker genes.

The identification of doublets in single cell transcriptome data remains computationally challenging. The interactive nature of CellView aids in cell doublet identification. In the PBMC data, ‘Subcluster-analysis’ reveals a mixture of lymphoid and myeloid gene expression within cluster 7 suggesting this cluster consists of doublets. Cluster 7 represents 2.3% of all cells in this data set, reflecting the number of expected doublets for the quantity of cells processed in this experiment. Thus, CellView can be utilized as a tool to pre-process of single cell data and remove doublets prior to final visualization.

The ‘Subcluster-analysis’ tab also provides a powerful tool to identify different cell types within clusters (where trade-offs between sensitivity and specificity in the chosen clustering algorithm may be insufficient to identify unique clusters) or a continuum of states within a cell type. For example, blood monocytes span a continuum of classical, intermediate, and non-classical subtypes in flow cytometry analysis of cell surface markers CD14 and CD16^11^. Two populations of cells within a cluster can be selected by square brush strokes for differential gene expression analyses (using a likelihood test) to identify biologically informative markers. For example, monocytes occupying cluster 4 in the PBMC data appear to contain two lobes (Fig.2b). Differential expression between these two lobes using the ‘Subcluster-analysis’ tool identified *CD16/FCGR3A* as the most differentially expressed gene marking the smaller lobe. This lobe also contained higher expression of MHC class II genes, an additional feature of non-classical blood monocytes. *CD14* is among the top 10 up-regulated genes in the large lobe, which include other classical blood monocyte markers (e.g. *S100A8, S100A9, S100A12*). Thus, this blood monocyte continuum defined by two cell surface molecules is detected by this transcriptome cytometry approach and represented by 837 parameters (*i.e*. genes) per cell. CellView data visualization may enable immunologists to explore further underlying biology within the blood monocyte compartment, such as investigating a subset of cells within the intermediate sub-cluster expressing *C1QA, C1QB*, and *C1QC*, markers of macrophage in tissue.

Dendritic cells (DCs) occupy clusters 6 and 8 in the PBMC data. Cluster 6 represents plasmacytoid DCs, expressing *CLEC4C/CD303, CD68, IL3RA/CD123* and *LILRA4/CD85g*. Myeloid DCs comprise cluster 8. CellView’s ‘Subcluster-analysis’ tool enables identification of both the common CD1C+ DC (Fig. 2c; 113/215 cells expressing *CLEC10A, CD1C)* and less adundant CD141+ DCs (Fig. 2d; 12/215 cells expressing *CLEC9A, IRF8)*. An additional layer of data we include in our .rds files are the genes and unique molecular identifiers (UMIs) detected per cell; this can enable identification of cell type biological features since RNA abundance (and therefore UMI count) often correlates with cell size^12^. Notably, non-classical blood monocytes and myeloid dendritic cells have the greatest numbers of UMIs detected per cell, at 3,719 and 5,645 respectively. In contrast, remaining cells have 2,305 UMIs per cell. Myeloid DCs are not noticeably larger than other PBMCs and non-classical blood monocytes are somewhat smaller in size than classical blood monocytes^11^ suggesting the RNA content is reflective of an underlying biological feature of these cells rather than cell size and may reflect the precursor relationship between non-classical monocytes and myeloid dendritic cells^13^.

We next applied CellView to human pancreatic islet single cell transcriptome data we generated on the Chromium system from a nondiabetic normal donor, which resulted in 4,806 cells sequenced to 87.1% saturation and detecting, on average, 1,848 genes and 7,686 molecules detected per cell. Our pipeline identified eight distinct clusters (Fig 1e). Using CellView’s ‘AllClusters’, and marker genes we had previously used to cell type in human islets^14^, we identified endocrine alpha, beta, delta, and gamma cell clusters and exocrine acinar, ductal, and stellate cell clusters. An 8th cluster represented endothelial cells (Fig 1f). Visual inspection of the 3D scatter plot displaying cells in tSNE space indicated two sub-clusters within the defined stellate cell cloud. The ‘Sub-cluster’ tool revealed, in addition to the stellate cells, a sub-cluster expressing the pericyte marker *RGS5*^*15*^. The close proximity of stellate cells and pericytes are likely a result of their shared mesenchymal origin, as both express *COL1A1* and *ACTA2*. Visual inspection of the ductal cell cluster identified a spread of cells suggestive of a continuum of cell states. Differential expression between cells at opposing ends of this continuum using the ‘Sub-cluster’ tool identified biologically meaningful differences. While all cells expressed *KRT19*, there was a transition from a *REG1A/AMBP*-positive (Fig 1g) to a *TFF1/TFF2/TFF3/FGF19/CAECAM6*-positive (Fig 1h) population. Whether these represent different spatially localized populations of epithelial cells within the pancreatic duct or different states of activation remains to be determined, but further highlights the utility of CellView to uncover putative novel biology.

These examples illustrate how CellView provides a powerful complement to current command line approaches to cluster and identify cell types in single cell experiments. This intuitive web application enables collaboration between biologists and computational analysts and increases the value of each single cell dataset. Moreover, the CellView framework provides a useful format to present these data in an interactive manner and can be broadly applied to single cell and bulk genomics assays with count matrix and cluster information. Until a complete atlas of cell-type transcriptomes has been defined, where a reference-based approach may prove more powerful for clustering and cell type identification^16^, CellView provides a useful tool to explore and characterize single cell data.

## METHODS

*Single cell RNA-seq* - PBMCs were purchased from AllCells, thawed quickly at 37°C and into DMEM supplemented with 10% FBS. Cells were quickly spun down at 400g, for 10min. Cells were washed once with 1 x PBS supplemented with 0.04% BSA and finally resuspended in 1 × PBS with 0.04% BSA. Viability was determined using trypan blue staining and measured on a Countess FL II. Briefly, 12000 cells were loaded for capture onto the Chromium System using the v2 single cell reagent kit (10X Genomics). Following capture and lysis, cDNA was synthesized and amplified (12 cycles) as per manufacturer’s protocol (10X Genomics). The amplified cDNA was used to construct an Illumina sequencing library and sequenced on a single lane of a HiSeq 4000.

Human islets from one nondiabetic deceased organ donor (UNOS ID ADIW417) were purchased from ProdoLabs and processed to obtain a single cell suspension as previously described^14^. Briefly, islets were dissociated using Accutase and filtered through a prewet cell strainer (BD) to collect single cells. The single cell suspension was prepared and loaded onto the Chromium System as described above.

*FASTQ generation and Alignments* - Illumina basecall files (*.bcl) were converted to fastqs using cellranger v1.3, which uses bcl2fastq v2.17.1.14. FASTQ files were then aligned to hg19 genome and transcriptome using the cellranger v1.3 pipeline, which generates a *gene vs cell* expression matrix. *Clustering and marker gene identification* - Cells with less than 500 total unique transcripts were removed prior to downstream analysis. Genes for clustering were selected based on normalized dispersion analysis. Cells were clustered using Barnes Hut t-SNE^9^ with the 1000 most over dispersed genes and clusters identified using DBSCAN (eps = 5.0, minpts=15). Differential gene expression was computed using edgeR^17^ and signature genes defined as genes upregulated 2 fold and FDR < 0.01 in all pairwise comparisons.

*Datasets and visualization* – Access to CellView from: https://www.jax.org/CellOmics

## ACKNOWLEDGEMENTS

We thank Karolina Palucka for helpful discussions. This work was supported by grants from NIAMS 1P50AR070594-01 (to PR), NCI 5P30CA034196-31 (PR) and The Jackson Laboratory scientific services core budget (MB, PR). Generation and analysis of the islet 10x Genomics dataset was supported by the Assistant Secretary of Defense for Health Affairs, through the Peer Reviewed Medical Research Program under Award No. W81XWH-16-1-0130 (to MLS). Opinions, interpretations, conclusions, and recommendations are solely the responsibility of the authors and are not necessarily endorsed by the Department of Defense.

## CONFLICT OF INTEREST

The authors declare no competing financial interests.

## CONTRIBUTIONS

MTB and PR conceptualized the software package. MTB wrote the software package. MTB analyzed the PBMC and Pancreatic islets data. MTB, PR and MS interpreted the data to identify cell types. MTB, PR and MS wrote the manuscript.

